# Haplotype Threading: Accurate Polyploid Phasing from Long Reads

**DOI:** 10.1101/2020.02.04.933523

**Authors:** Sven D. Schrinner, Rebecca Serra Mari, Jana Ebler, Mikko Rautiainen, Lancelot Seillier, Julia J. Reimer, Björn Usadel, Tobias Marschall, Gunnar W. Klau

**Affiliations:** Algorithmic Bioinformatics, Heinrich Heine University Düsseldorf, Universitätsstr. 1, 40225 Düsseldorf, Germany; Center for Bioinformatics, Saarland University, 66123 Saarbrücken, Germany; Graduate School of Computer Science, 66123 Saarbrücken, Germany; Forschungszentrum Jülich IBG-4, Wilhelm- Johnen-Str., 52428 Jülich, Germany; Institute for Biology I, RWTH Aachen, Worringer Weg 3, 52078 Aachen, Germany; Max Planck Institute for Informatics, 66123 Saarbrücken, Germany; Cluster of Excellence on Plant Sciences (CEPLAS), Heinrich Heine University Düsseldorf, Universitätsstr. 1, 40225 Düsseldorf, Germany

**Keywords:** polyploidy, phasing, haplotypes, cluster editing, high-throughput nucleotide sequencing, plant science, sequence analysis

## Abstract

Resolving genomes at haplotype level is crucial for understanding the evolutionary history of polyploid species and for designing advanced breeding strategies. As a highly complex computational problem, polyploid phasing still presents considerable challenges, especially in regions of collapsing haplotypes.

We present WhatsHap polyphase, a novel two-stage approach that addresses these challenges by (i) clustering reads using a position-dependent scoring function and (ii) threading the haplotypes through the clusters by dynamic programming. We demonstrate on a simulated data set that this results in accurate haplotypes with switch error rates that are around three times lower than those obtainable by the current state-of-the-art and even around seven times lower in regions of collapsing haplotypes. Using a real data set comprising long and short read tetraploid potato sequencing data we show that WhatsHap polyphase is able to phase the majority of the potato genes after error correction, which enables the assembly of local genomic regions of interest at haplotype level. Our algorithm is implemented as part of the widely used open source tool WhatsHap and ready to be included in production settings.

## Background

Polyploid genomes have more than two homologous sets of chromosomes. Polyploidy is common to many plant species, including important food crops like potato (*Solanum tuberosum*), bread wheat (*Triticum aestivum*) and durum wheat (*Triticum durum*). Resolving polyploid genomes at the haplotype level, i.e., assembling the sequences of alleles residing on the same chromosome, is crucial for understanding the evolutionary history of polyploid species: Evolutionary events, such as whole genome duplications, can be traced back and reveal the ancestry of polyploid organisms [1]. Beyond that, knowledge of haplotypes is key for advanced breeding strategies or genome engineering, especially for improving yield quality in important crop species [1, 2, 3].

In this work we focus on phasing from long read information. Plant genomes typically exhibit many highly repetitive regions and frequently underwent structural variation events, rendering alignments from short reads alone problematic. Although long reads suffer from a higher number of sequencing errors, they align better to the reference genome and span more variant positions. Consequently, there are larger overlaps between read pairs, which is the key information for molecular phasing methods. This is especially important for polyploid phasing, where the assignment must distinguish not only between two but between *k* haplotypes.

While phasing diploid genomes using long reads has become a routine step, polyploid phasing still presents considerable challenges [4]. Higher ploidy increases the complexity of the underlying computational problem: In the diploid case, assembling one haplotype over all heterozygous variants directly determines the complementary second haplotype. For genomes of higher ploidy, this is not the case. In addition, polyploid genomes usually exhibit larger regions of two or more identical haplotypes. The Minimum Error Correction (MEC) model [5], which is the most common and successful formalization for diploid haplotype assembly from sequence reads, is not suited to distinguish between locally identical haplotypes. It aims to minimize the number of corrections that are applied to the reads in order to partition them into distinct sets such that reads from the same partition belong to the same haplotype. The MEC score is the minimum number of necessary corrections. In the MEC model it does not pay off to assign identical haplotypes. Hence, in regions of locally similar haplotypes, this model is likely to result in incorrect haplotype assignments, see Figure 1. Consequently, MEC-based approaches for polyploid phasing struggle in such regions and, beyond that, face the challenge that dynamic programming techniques for MEC [6] quickly become infeasible in practice.

**Figure 1:**
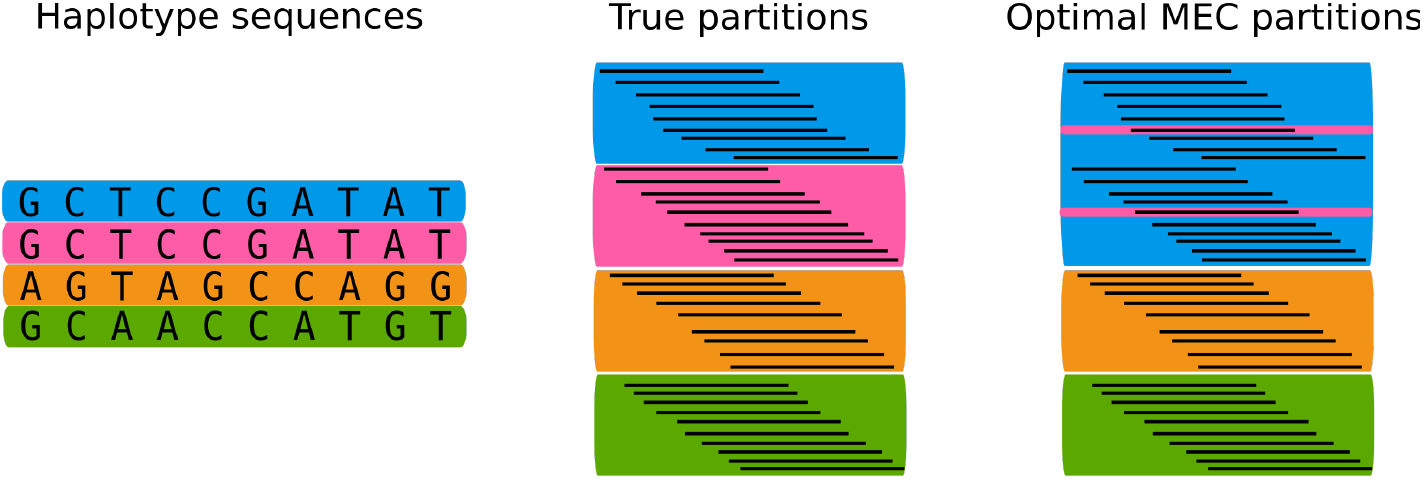
MEC model in collapsed regions. Frequently, in polyploid organisms, two or more haplotypes are locally identical on larger stretches of heterozygous sites, as shown by the pink and blue haplotype sequences in the left picture. The MEC model favors assigning the reads of two haplotypes to only one partition, because the spare partition can be used to collect noisy reads, which gives a lower MEC score but also results in unbalanced and likely wrong partitions.

### Related work

Throughout the last years, a few polyploid phasing methods have already been proposed. In 2013, Aguiar et al. were the first to introduce a theoretical framework for polyploid haplotype assembly with the HapCompass [7, 8] model, which is based on spanning trees and uses the Minimum Weighted Edge Removal (MWER) criterion. In 2014, Berger et al. introduced HapTree [9], a maximum likelihood approach to discover the most likely haplotypes given aligned read data. To address the problem of computational complexity, the most likely haplotypes are assembled for a small set of SNP positions first and then iteratively extended, keeping only the most likely sub-solutions in each step. HapTree was shown to outperform HapCompass in terms of accuracy and runtime [9, 10]. Together with SDhaP [11], a semi-definite programming approach based on an approximate MEC criterion, HapCompass and HapTree were evaluated and compared to each other in a simulation study conducted by Motazedi et al. [10] in 2017. The study, where simulated data of the tetraploid potato genome as model organism was used, revealed that, out of the compared methods, HapTree provided the best results in terms of precision. However, it also showed the highest time and memory requirements and often suffered from low recall. SDhaP showed low performance in regions of locally similar haplotypes, which is probably related to the underlying MEC model. For ploidies above six, HapCompass was the only implementation to remain stable, although it showed an overall poor performance. As a result, none of the methods came out to be applicable for practical use due to computational inefficiency that prohibits scaling to large genomic regions as well as frequent failures and low overall accuracy. In fact, the authors conclude that there is “clearly room for improvement in polyploid haplotyping algorithms” [10].

H-PoP [12] was shown to outperform these previous approaches both in accuracy and runtime and is since then considered as the state-of-the-art method. It consists of a model called Polyploid Balanced Optimal Partition (PBOP) that aims to create *k* partitions of sequence reads that minimize two measures: Reads from one partition are supposed to be equal on as many variant loci as possible, whereas reads from different partitions should contain as many differences as possible. For *k* = 2, this equals the diploid MEC model and can thus be seen as a polyploid generalization of MEC. When genotype information is present, genotype constraints are added to the model; the appropriate extension is then referred to as H-PoPG.

More recent advances have not proven to be useful for whole-genome single-individual haplotyping, like PolyHarsh [13], a Gibbs sampling method that is also based on the MEC model and has only been shown to work on very small artificial examples, TriPoly [14] that infers haplotypes from trio data and thus requires family data, and SDA [15]. The latter provides two algorithms based on a discrete matrix completion approach and correlation clustering, respectively, and is used to resolve segmental duplications of higher ploidy during genome assembly. However, it is not designed to scale to the whole genome.

Other matrix-based models are SCGD-hap [16], a structurally constrained gradient descent approach, and AltHap [17], which builds on SCGD-hap and aims to solve an iterative sparse tensor decomposition problem. This model yielded results similar to those of H-PoP, but also relies on MEC.

Some tools have been proposed that do not work well with long read data. The work by [18] is based on minimum fragment removal. The long and relatively erroneous long reads would lead to a removal of too much data. RANBOW [1] uses allele co-occurrences on small sets of sampled positions in overlapping short reads. This approach is susceptible to high error rates found in long reads, as it seeds the phasing on local partitions of reads based on their allele combination on the small position samples. Thus, a large portion of the reads are clustered incorrectly and a lot of overlapping position samples are required to correct these mistakes.

Apart from the limitations of the underlying model, current methods do not give reliable information about the accuracy of the resulting haplotypes since these are either output in one consecutive sequence or in very long blocks. In particular, this means that there is no information about the positions of likely switch errors. Thus, large regions of the resulting haplotypes might be out of place, but it is not possible to identify these regions, which makes the results very difficult to use in practice. In the H-PoPG algorithm, for example, the haplotypes are only divided into blocks if there is no read that covers the affected neighboring variant loci. Further uncertainties in the phasing are not considered in the model and thus not reported.

### Contribution

To our knowledge, there is currently no method that is designed to properly address polyploid phasing by offering an accurate model and is able to produce reliable blocks according to the phasing certainty while at the same time being computationally efficient and thus applicable in practice. To address this gap, we present WhatsHap polyphase, a method that departs from the MEC model in order to deal with the additional challenges arising in polyploid phasing. By taking coverage into account via a newly established threading step, WhatsHap polyphase is able to detect and properly phase regions where multiple haplotypes coincide. Additionally, our method is able to integrate information from input genotypes for accurate phasing results.

We introduce cuts within the haplotypes at positions with increased phasing uncertainty and thereby output phased blocks that ensure high accuracy within the fragments. We provide a sensible way to compute these block boundaries at varying, user-defined degrees of strictness. This way, we enable a configurable trade-off between longer blocks that potentially contain errors and shorter but highly accurate blocks.

We demonstrate on a simulated data set that WhatsHap polyphase returns results that are around three times more accurate than those computed by the state-of-the-art tool H-PoPG, in particular in regions of identical haplotypes, where our method phases with around seven times lower switch error rates than the competition. The efficient implementation of WhatsHap polyphase allows for scaling to gigabase-sized genomes, while being sufficiently fast: an artificial human tetraploid Chr01 (249Mb) is phased in less than 3.5 hours on a single core of a standard desktop.

WhatsHap polyphase is ready to be included in production settings since it is implemented as part of the widely used open source tool WhatsHap (https://whatshap.readthedocs.io), offering convenient usage by supporting standard input and output format (BAM and VCF). We used the tool to phase real potato data, assign corrected long reads to haplotypes and to locally assemble reads. WhatsHap polyphase is available at https://bitbucket.org/whatshap/whatshap.

## Results

### Phasing Model and Algorithm

WhatsHap polyphase is a novel two-stage approach that produces accurate haplotypes for polyploid genomes using data from single-molecule sequencing technologies. See Figure 2 for an overview of the method.

**Figure 2:**
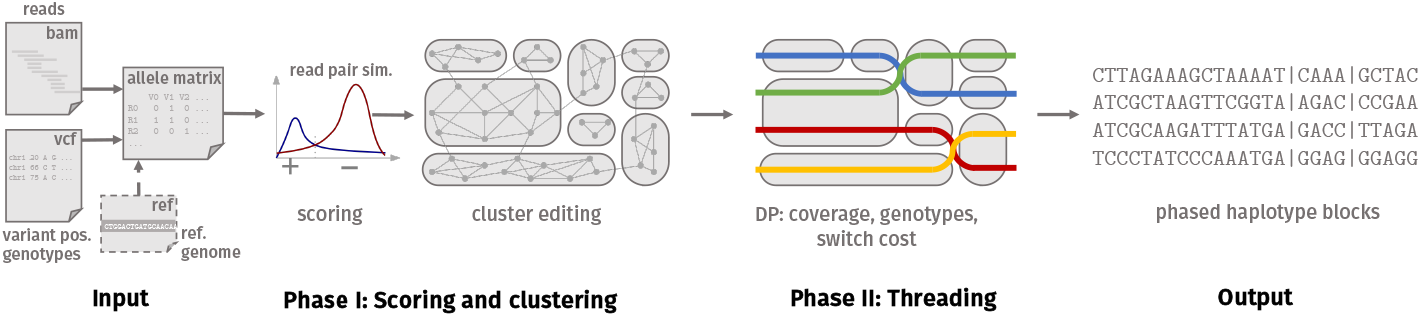
Overview of WhatsHap polyphase. The input allele matrix results from a given BAM and VCF file and an optional realignment step. Phase I: Statistical scoring of each read pair classifies them into belong to the same or to different haplotypes. The scores are used as weights for a graph over all reads, which is clustered by Cluster Editing (grey round shapes). Phase II threads *k* haplotypes (colored lines) through the clusters (here *k* = 4) balancing coverage violations and switch costs while respecting the genotype information. This results in *k* phased haplotypes, subdivided into blocks (vertical lines).

The first phase of the algorithm uses cluster editing [19] to find clusters of reads which are likely to originate from identical haplotypes. In short, this is done by computing a statistical similarity score for each pair of reads and constructing a graph using the reads as nodes and the scores as edge weights [20]. The size of the graph makes it infeasible to solve cluster editing to optimality in reasonable time, so we rely on an iterative heuristic to produce accurate clusters. We deliberately make no assumptions on the ploidy at the clustering stage. In particular, reads of multiple haplotypes that are locally identical end up in the same cluster.

The second phase consists of the actual haplotype assembly by threading *k* haplotypes through the set of clusters obtained in the first phase. We take the position-wise read coverage of each cluster into account to determine the number of haplotypes threaded through each cluster. In contrast to MEC-based models, this allows us to handle genomic regions where some haplotypes are locally identical by allowing that multiple haplotypes run through the same cluster. During the threading step, we further expect haplotypes to stay in the same cluster for as long as possible and ensure that the consensus genotype fits the input genotype, if provided. We cut the phasing into blocks at variant pairs showing insufficient phasing confidence to increase its accuracy at the cost of decreased phasing block lengths. See the Methods section for details of WhatsHap polyphase.

### WhatsHap polyphase produces accurate results

To demonstrate that WhatsHap polyphase works well in practice, we ran it on an artificial tetraploid dataset at different coverages and compared our results to those of H-PoPG, the state-of-the-art method for polyploid phasing. We used common evaluation statistics that capture different properties of haplotype sequences to compare the solutions computed by both tools to ground truth haplotypes available for our data sets.

#### Evaluation statistics

For ploidy *k*, a set of ground truth haplotype sequences *h* = {*h*_1_,…, *h_k_*} and predicted haplotypes 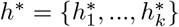, we compute the number of Hamming errors HE as

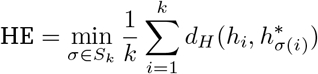

where *S_k_* represents the permutation group on {1,…, *k*} and *d_H_*() the Hamming distance between two sequences. The *Hamming rate* (*HR*) is then defined as the sum of Hamming errors divided by the total number of all phased variants. If subtracted from 1, the Hamming rate is equivalent to the *reconstruction rate* and the *correct phasing rate* presented in [14] and [12], respectively.

A well established evaluation metric for diploid phasing is the *switch error rate (SER),* for which we use a polyploid version. Instead of counting the number of incorrect alleles on each haplotype, the SER counts the minimum number of switches, i.e., how often the assignment between predicted and true haplotypes must be changed in order to reconstruct the true haplotypes from the predicted ones. The polyploid extension of the switch error was already introduced as the *vector error rate* in [9].

More formally, for every position *j* let Π_*j*_ be the set of one-to-one mappings between *h* and *h*\*, such that for each *π* ∈ Π_*j*_ it holds that 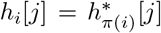 for all haplotypes *h_i_*. The switch error rate is then defined as:

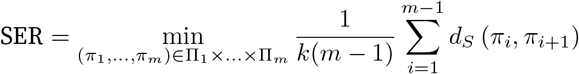

where *m* is the number of variants and *d_S_* (*π_i_, π*_*i*+1_) the number of different mappings between *π_i_* and *π*_*i*+1_.

If the genotype of *h*\* is not equal to the genotype of *h* for every position, the set Π_1_ × … × Π_*m*_ is empty and the vector error cannot be computed. Therefore we compare only those positions, on which the predicted genotype is correct and report the fraction of *missing variants (MV),* that is, either unphased or incorrectly genotyped variants, separately.

Phasing tools may not phase the entire input region as one set of haplotypes. If the phasing between two consecutive variants is too uncertain (e.g., if not enough reads cover both variants), the phasing might be split into *blocks*. In our evaluation, we applied the HR and SER on all reported phasing blocks separately and aggregated them. More precisely, we summed up the number of respective errors and divided them by the total number of variants (HR) or by the total number of variants excluding the first variant in every block (SER). Since this favors shorter blocks, we also included the N50 block length into our evaluation, which is the smallest block length needed to cover 50% of the considered genomic region when using only blocks of that size and larger.

#### Testing on artificial tetraploid human

We generated a tetraploid version of human chromosome 1 by combining sequencing data of two individuals (NA19240 and HG00514), for which high-quality trio-based haplotype information is available [21]. We refer to these haplotypes as ground truth haplotypes. We merged PacBio sequencing data for these two samples to produce tetraploid data at different coverages (40 × and 80 ×). Using the read simulator PBSIM [22], we additionally generated equivalent simulated data sets with known read origin.

We ran WhatsHap polyphase and H-PoPG and compared the resulting phasings to the ground truth haplotypes. H-PoPG defines phased blocks based on the connected components of the underlying reads by introducing cuts between pairs of variants not connected by any sequencing reads. Per default, WhatsHap polyphase uses a more sensitive approach (see Methods section) typically leading to shorter but more accurate haplotype blocks. Additionally, our algorithm supports different levels of block cut sensitivities, which allow to balance block length against block accuracy. In order to provide a better comparison of both tools, we ran WhatsHap polyphase with different configurations, which can be seen in Figure 3.

**Figure 3:**
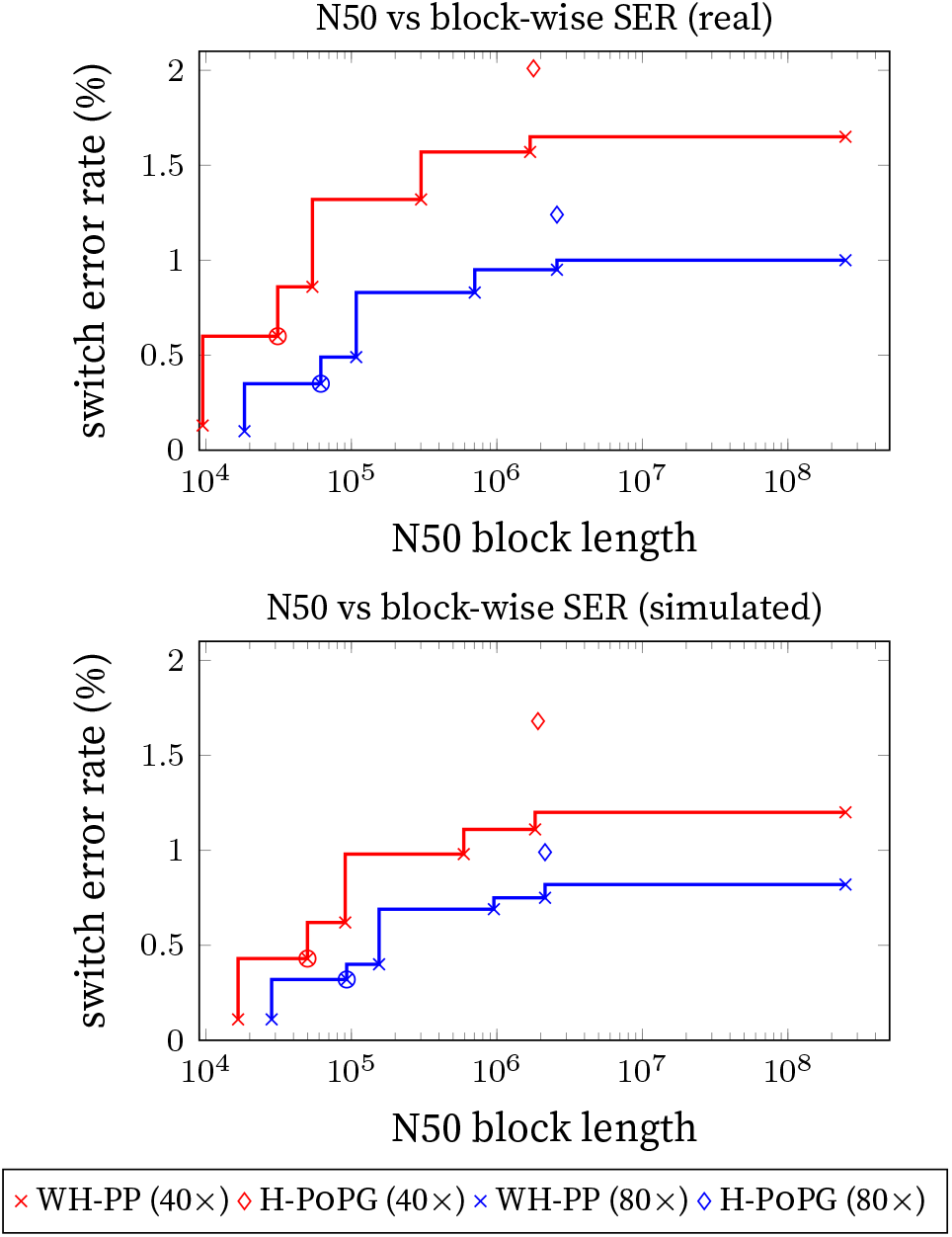
N50 block lengths and the respective block-wise switch error rates for different block cut strategies of WhatsHap polyphase (default strategy marked by a circle) on the real read dataset (top) and the simulated dataset (bottom) with 40 × and 80 × coverage.

Even when forcing our tool to yield block lengths as computed in H-PoPG we observe around 25% lower switch error rates among the tested datasets (Figure 3, see Suppl. Figure 7 for Hamming error rates). As expected, higher coverage has a positive effect on the error rates. More sensitive block cuts, and in particular the default setting for WhatsHap polyphase, lead to a dramatic decrease in switch error rates.

Table 1 shows all used evaluation metrics on H-PoPG and WhatsHap polyphase for their default settings. We can see that WhatsHap polyphase phases more accurately, with at least three times lower switch error rates than H-PoPG on the varying data sets. For the Hamming rate the differences are even larger. Among other reasons, this is caused by the block cut policy of H-PoPG, leading to switch errors on sparsely connected variants, which have a big impact on the global correctness of the phasings. The configurable block cut strategy of WhatsHap polyphase allows to maintain accurate blocks with low Hamming rates.

**Table 1:** Comparison of WhatsHap polyphase and H-PoPG on real (a) and simulated (b) datasets. Performances are based on the switch error rate (SER), block-wise Hamming rate (HR) and N50 for the block size. The total length of the chromosome is 249 Mb.

| coverage | method | SER (%) | HR (%) | N50 (bp) | runtime (s) |
| --- | --- | --- | --- | --- | --- |
| 40× | WH-PP | 0.60 | 1.57 | 31140 | 2909 |
|  | H-POPg | 2.01 | 27.53 | 1785293 | 1853 |
| | $SER(\frac{H-POPg}{WH-PP})$ | 3.35 | | | |
| 80× | WH-PP | 0.35 | 2.01 | 64315 | 11559 |
|  | H-POPg | 1.24 | 27.66 | 2587104 | 3668 |
| | $SER(\frac{H-POPg}{WH-PP})$ | 3.54 | | | |

(a) real read data
| coverage | method | SER (%) | HR (%) | N50 (bp) | runtime (s) |
| --- | --- | --- | --- | --- | --- |
| 40× | WH-PP | 0.43 | 1.72 | 51416 | 1792 |
|  | H-POPg | 1.68 | 26.38 | 1917094 | 1215 |
| | $SER(\frac{H-POPg}{WH-PP})$ | 3.91 | | | |
| 80× | WH-PP | 0.32 | 3.39 | 103287 | 5260 |
|  | H-POPg | 0.99 | 25.65 | 2142893 | 2447 |
| | $SER(\frac{H-POPg}{WH-PP})$ | 3.09 | | | |

(b) simulated read data

While comparing the WhatsHap polyphase phasing against the ground truth, we noticed that a small portion of variants was marked as unphased, namely 0.31% and 0.07% for the real reads with 40 × and 80 × coverage, respectively. This happens when a variant is supposed to be heterozygous, but is reported as homozygous by WhatsHap polyphase due to the actual alleles of the reads. H-PoPG strictly sticks to the input genotypes and never deviates from them, resulting in no unphased variants. However, in practice it is not necessarily a mistake to deviate, as the given genotype could be wrongly called by another tool. For the simulated reads the fraction of unphased reads by WhatsHap polyphase goes down to 0.19% and 0.01% (for 40 × and 80× coverage), indicating that for real data the genotypes are more likely to contradict the observed read alleles.

### Identifying collapsing regions

We define regions in the genome where two or more haplotypes share the same sequence for at least 50 variant positions as *collapsing regions.* For MEC-based approaches, these parts are particularly difficult to phase since different configurations of haplotypes with locally identical sequences are not distinguishable based on their MEC scores and the MEC model exploits this to explain sequencing errors with “noise” haplotypes.

We evaluated the ability to correctly assemble haplotypes in these regions. Again, both WhatsHap polyphase and H-PoPG were run on chromosome 1 of the simulated and real datasets with 40 × and 80 × coverage, respectively. Collapsing regions take up a large part (17.28%) of the simulated Chr01.

Table 2a shows the results. It can be seen that the differences between switch error rates achieved by H-PoPG and by WhatsHap polyphase are remarkably higher in the case of collapsing regions than for the rest of the genome. In comparison to WhatsHap polyphase, the switch error rate of H-PoPG is around 7 times higher in collapsing regions, while on average throughout the whole chromosome, this factor is only 3.37. For higher coverage, these values are further increased to 7.5 and 3.5, respectively. The closest results are achieved in non-collapsing regions, i.e., regions where either all haplotype sequences are unique or coincide on fragments shorter than 50 variants. In these regions, H-PoPG returns 3.13 times more switch errors.

**Table 2:** Comparison between the resulting switch error rates of WhatsHap polyphase (WH-PP) and H-PoPG on collapsing regions over at least 50 variants as compared to noncollapsing regions and the average throughout the genome. Results (switch error rates in %) are presented for Chr01 of the real (a) and simulated (b) dataset on both 40 × and 80 × coverage. The third row marks the quotient between the switch error rate of H-PoPG and that of WhatsHap polyphase to highlight by which magnitude the results differ.

| coverage | method | collapsing regions | non-collapsing regions | total |
| --- | --- | --- | --- | --- |
| 40× | WH-PP | 0.29 | 0.69 | 0.60 |
|  | H-POPG | 2.02 | 2.16 | 2.02 |
| | $SER(\frac{H-POPG}{WH-PP})$ | 6.97 | 3.13 | 3.37 |
| 80× | WH-PP | 0.14 | 0.46 | 0.35 |
|  | H-POPG | 1.05 | 1.30 | 1.24 |
| | $SER(\frac{H-POPG}{WH-PP})$ | 7.50 | 2.83 | 3.54 |

(a) real read data
| coverage | method | collapsing regions | non-collapsing regions | total |
| --- | --- | --- | --- | --- |
| 40× | WH-PP | 0.18 | 0.45 | 0.43 |
|  | H-POPG | 2.01 | 1.63 | 1.68 |
| | $SER(\frac{H-POPG}{WH-PP})$ | 11.17 | 3.62 | 3.91 |
| 80× | WH-PP | 0.08 | 0.37 | 0.32 |
|  | H-POPG | 0.94 | 0.98 | 0.99 |
| | $SER(\frac{H-POPG}{WH-PP})$ | 11.75 | 2.65 | 3.09 |

(b) simulated read data

For the simulated data (see Table 2b), the differences are even more striking, especially on 80 × coverage. In regions with coinciding haplotypes, WhatsHap polyphase outperforms H-PoPG by a factor of up to 11.75. Compared to the average quotient of 3.09, WhatsHap polyphase thereby yields an almost 4 times higher reduction in switch error rates in collapsing regions. On lower coverage, similar results are obtained.

As for the previous experiments, we repeated this analysis with block lengths computed as in H-PoPG. The results of this second run are presented in Suppl. Table 3.

### Potato data

We applied our algorithm to real sequencing data for tetraploid potato (*Solanum tuberosum*), for which we generated paired-end short Illumina and long Oxford Nanopore reads. In a first step, we aligned the reads produced by the different technologies to the potato reference genome published by the Potato Genome Sequencing Consortium (PGSC) [24]. We observed unbalanced coverage distributions for the alignments, especially for the short Illumina reads, hinting towards a high number of structural variations and rearrangements being present in the data (Figure 4a). Thus, the Illumina reads are ill-suited for reliable variant calling as their short length makes it more difficult to unambiguously align them to the reference.

**Figure 4:**
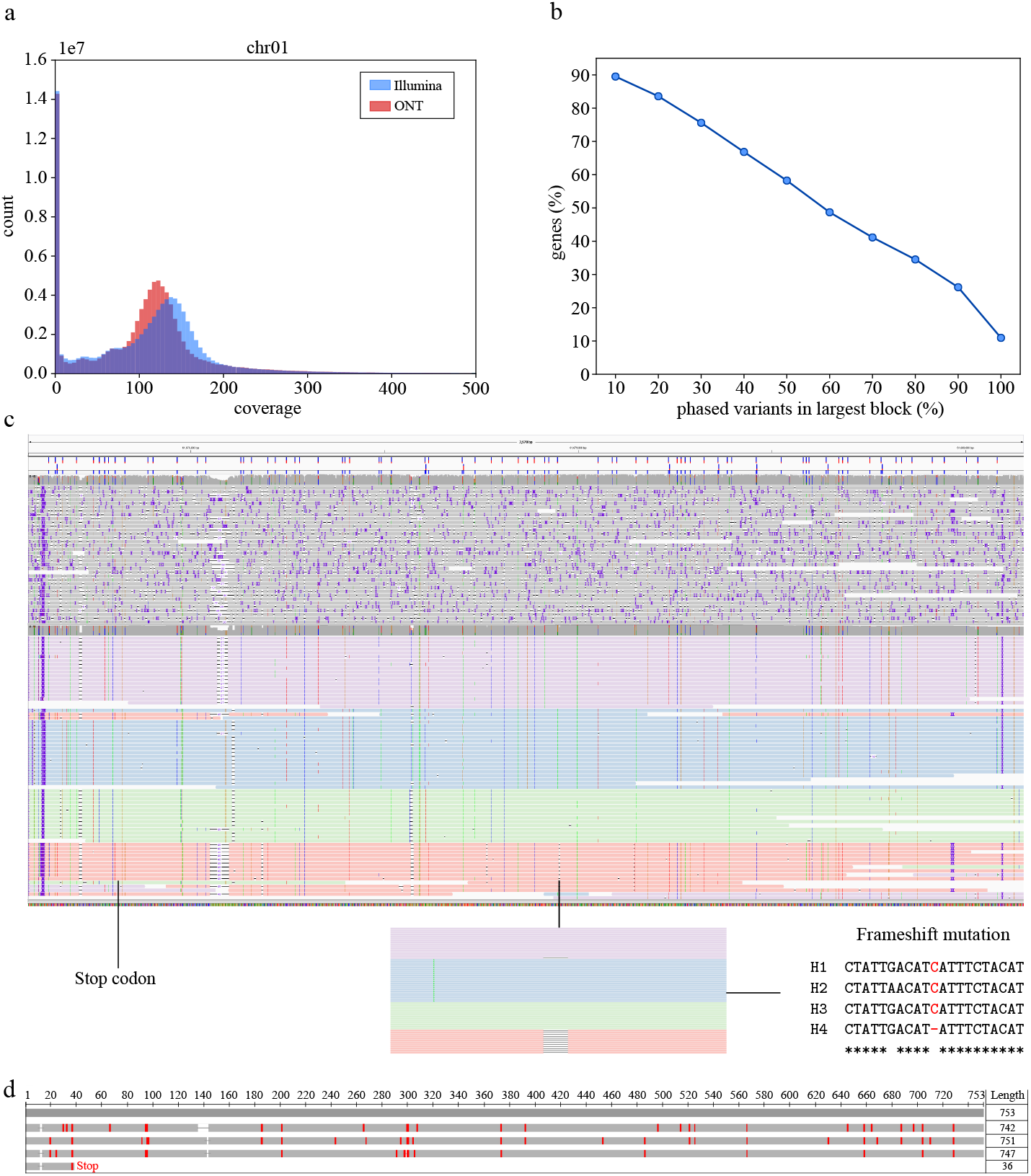
Phasing of potato genome. (**a**) Per-base coverage distribution of Illumina and ONT MinION alignments on Chr01. (**b**) For each gene, we consider only the longest phased block. The x-axis shows how many of the heterozygous variants were at least phased in the largest block, the y-axis displays for how many genes this was the case. (**c**) IGV [23] screenshot showing alignments of uncorrected (top) and corrected MinION reads (bottom) of F gene on Chr04. The corrected reads are colored (red, green, blue, purple) according to the haplotypes WhatsHap polyphase assigned them to. (**d**) Multiple sequence alignment of the ORFs detected in the four haplotype sequences. The uppermost gray sequence represents the reference, the others correspond to the four haplotypes (same order as in panel c)

We therefore relied on the much longer nanopore reads to identify SNPs that we can later use for phasing. However, Oxford Nanopore reads typically come with high sequencing error rates, complicating the calling process. In order to obtain reliable variant positions and genotypes from these error-prone reads, we ran an error correction pipeline [25] to reduce the number of sequencing errors (see Methods section). Figure 4c shows an exemplary IGV [23] screenshot of uncorrected reads (top) and corrected (bottom) for the FRIGIDA-like protein 5 isoform X2 gene. Next, we ran minimap2 [26] to align the corrected nanopore reads to the potato reference genome and called variants using FreeBayes [27]. To verify the genotypes produced in this way, we added an additional step to WhatsHap polyphase that re-genotypes the positions based on the nanopore reads prior to phasing and only keeps those variants, for which the newly predicted genotype matches the one reported by FreeBayes (see Methods section).

We focused on the potato genes [24] as they are biologically interesting for phasing. Of in total 36274 genes containing heterozygous variants after calling and retyping, 91% could be (at least partially) phased by WhatsHap polyphase. On average, about 2.13 phased blocks were produced per gene. Furthermore, for each gene, we took the longest phased block and determined the number of phased variants inside of this block relative to the total number of heterozygous variants reported in the gene. For 58% of all genes, the longest block covered at least half of all variants within the respective genomic interval (see Figure 4b). For about 11% of all genes, the longest block covered all heterozygous variants. These are genes that we can phase completely in a single block. The fraction of genes for which the longest block covered at least 90% of all heterozygous variants is about 26%.

We used the FRIGIDA-like protein 5 isoform X2 (accession: XP_015169713) gene as an example to demonstrate how WhatsHap polyphase enables haplotype-resolved assembly. We extracted the phasing of the longest phasing block reported for this gene and separated the reads by haplotype. In order to do so, we extended the commands whatshap haplotag and whatshap split, previously implemented in the diploid version of whatshap, to higher ploidies. Briefly, the idea is to tag each sequencing read according to the computed haplotype sequence it is most similar to and separate the reads based on these tags (see Methods section). The reads shown in Figure 4c are colored according to the resulting haplotype assignments. In the next step, we separately ran wtdbg2 [28] on each haplotype-specific read set to produce local assemblies of the four haplotypes. Suppl. Figure 8 shows a visualization of a multiple sequence alignment of these haplotypes. We ran the NCBI ORFfinder [29] on each of the assemblies and detected a long ORF in the first three haplotypes representing the FRIGIDA like coding sequence. For the fourth haplotype we could not detect a corresponding ORF, as the putative FRIGIDA gene in the fourth phase showed an early STOP codon highlighted in Figure 4c. Interestingly, the fourth phase showed an additional frameshift mutation shown in the inset of Figure 4c where only the phasing information provides the information that this is linked to the premature STOP codon highlighting the necessity of (local) phasing to understand gene architecture. Using COBALT [30], we generated multiple sequence alignments of the amino acid sequences resulting from these three ORFs and the corresponding reference sequence (Figure 4d). The three sequences show an overall good alignment with the reference with small differences, which may serve as an input for functional follow-up studies.

### Runtimes

We show the runtimes of WhatsHap polyphase and H-PoPG for phasing the artificial human Chr01 in Table 1. Both programs were run on a single core on a dual socket machine (2 × Intel Xeon E5-2670 v2) with 256GB of memory. At coverage 40×, WhatsHap polyphase took about 49 min to phase the real data, while H-PoPG took about 30 min. WhatsHap polyphase phased the simulated data set in about 30 min and H-PoPG in 20 min. At coverage 80×, WhatsHap polyphase took 3.2 hours on the real data and H-PoPG 1 h. On the simulated data, WhatsHap polyphase took 1.5 hours and H-PoPG 41 min for phasing.

## Discussion

We introduce a novel two-stage algorithm enabling accurate haplotype phasing of polyploid genomes. Our model consists of two phases performing a clustering of the reads based on their similarity and assembling the final haplotypes through the resulting clusters. We emphasize that unlike approaches based on solving the MEC problem, WhatsHap polyphase takes coverage of the read clusters into account to resolve regions with multiple coinciding haplotypes. Additionally the phasing can be cut at low confident positions to maximize phasing accuracy.

Applying our algorithm to chromosome 1 of a tetraploid dataset created of human samples HG00514 and NA19240 showed that in comparison with H-PoPG, the current state-of-the-art phasing method, WhatsHap polyphase returns around 61% and 68% lower switch error rates on real and simulated data at different coverages. The phased blocks produced using the default settings of WhatsHap polyphase are shorter compared to the ones reported by H-PoPG, but by a factor of three times more accurate. We offer a way to configure the trade-off between block length and accuracy via a parameter and are thus also able to compare both tools with the block sizes used by H-PoPG. In this case, block lengths are very similar, but WhatsHap polyphase still returns lower switch error rates. Note, however, that we usually recommend more conservative settings in order to obtain more interpretable results.

The Hamming rate behaves differently from the switch error rate, as pointed out in the result section. The reason for that is that most of the actual mistakes done by the phasing algorithms are in fact switch errors, where haplotype sequences are wrongly connected. The Hamming rate is very sensitive to these errors, because a single switch on two haplotypes in the middle of a block can potentially cause 50% of the variants being phased wrongly on the two affected haplotypes. While the additional block cuts eliminate only 50-65% of the switches, the impact on the Hamming rate is much higher. Moreover, if a sufficient number of switches is already present in the data, additional switches do not cause a substantial increase in the Hamming rate anymore, as newly introduced switches have a chance to cancel out old ones. Even though H-PoPG does outperform WhatsHap polyphase in terms of Hamming rates for equally long blocks, it is debatable how much the phasings benefit from this, as the phased blocks are very unreliable for both algorithms.

Furthermore, we show that the coverage-aware approach of haplotype threading is able to resolve regions where multiple haplotypes coincide, which occur frequently in polyploid genomes. A comparison to H-PoPG shows that WhatsHap polyphase performs particularly well in these regions. The switch error rates are 7 times higher in H-PoPG for the real data and more than 11 times higher for the simulated dataset. When using larger blocks according to the block definition of H-PoPG, the switch error rate of H-PoPG is still more than 3 times higher in these collapsing regions as opposed to 1.22 times, on average. Within chromosome 1 of our simulated dataset, with a total length of 249MB, we found around 17% of the genome to be part of long collapsing regions over at least 50 variants. These results clearly highlight the limitation of MEC-based approaches with regard to these regions and the need for phasing methods that address this problem.

Finally, we present a typical use case of polyploid phasing using real sequencing data of potato. Due to the high genomic diversity and lack of high quality reference sequences, large-scale polyploid phasing remains challenging. We restricted our analysis to the gene regions and use the FRIGIDA-like protein 5 isoform X2 gene as an example to demonstrate that our polyploid phasing tools enable haplotype-resolved assembly of polyploid organisms.

## Conclusions

Polyploid phasing is a difficult technological and computational problem. Current state-of-the-art tools rely on the Minimum Error Correction model, which is successful for diploid phasing, but has limitations in the conceptually and computationally far more complex polyploid case. We provide an implementation that departs from the MEC paradigm and instead uses a novel clustering and threading method, taking coverage and genotype information into account. Doing so, it represents the first algorithm designed to specifically handle locally identical haplotypes and, in consequence, performs significantly better in such regions than the state-of-the-art. To our knowledge, it is also the first approach that offers a configurable trade-off between the lengths of phased haplotype blocks and phasing accuracy to fit the user’s individual needs. Our implementation scales to whole genomes while being sufficiently fast.

Current challenges lie in resolving more switch error locations, as they either lead to block cuts or to switch errors, which have a high impact on the Hamming rate. Also, the running time of our approach scales exponentially with increasing ploidy, which requires further optimization to enable phasings with ploidy higher than six. Another limitation in practice is given by the fact that alignment-based phasing methods heavily depend on the quality of the alignments and the subsequent variant calls. In case of strong deviations from the reference genome, as, for example, in large regions of our proof-of-concept potato phasing study presented in this paper, any alignment-based method that relies on the reference genome will struggle.

On good quality reference genomes such as the artificial tetraploid benchmark genome proposed in this paper we show that our method WhatsHap polyphase delivers haplotype reconstructions with significantly lower error rates compared to the state-of-the-art tool H-PoPG. Our algorithm is implemented as part of the widely used open source tool WhatsHap and is hence ready to be included in production settings.

## Methods

Here, we present the phasing algorithm in detail. We denote with *k* the ploidy of the phased genome, with *n* the number of heterozygous variants in the genomic region of interest and with m the number of reads. We assume that all variants are biallelic, denoting the major allele with 0 and the minor allele with 1. Each read *r* is represented by a sequence *r*_0_,…, *r*_*n*–1_ of length *n* over the alphabet Σ = {0,1, –} such that *r_i_* is the allele for the *i*-th variant and “−” indicates an uncovered variant. We use olp(*r, s*) = |{*i* | *r_i_*, *s_i_* ∈ {0,1}}| to denote the size of the overlap (number of shared variants) between two reads *r, s* and dis(*r, s*) = |{*i* | *r_i_*, *s_i_* ∈ {0,1}, *r_i_* ≠ *s_i_*}| for the number of disagreements between *r* and *s*. The ratio between these values is a value between 0 and 1 and called the *Hamming rate* between two reads. The true (and to us unknown) haplotype of a read *r* is denoted as *H*(*r*) ∈ {0,…, *k* – 1}. The objective is to find *k* sequences 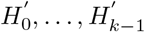 of length *n* over *Σ*, which are close or identical to the original haplotypes *H*_0_,…, *H*_*k*–1_.

### Clustering

The first step of our algorithm is to cluster reads that are likely to originate from the same haplotype. The clustering is based on pairwise similarity of overlapping reads. The similarity scores of the read pairs are then used in the clustering process. Two reads with an overlap of less than 2 variants are not considered as overlapping and always get a neutral score of 0.

We make two assumptions about the reads for the scoring scheme. First, all true haplotypes are expected to be equally frequent among the reads. Second, the Hamming rate between all pairs of haplotypes is expected to be the same (i.e., all haplotypes are equally different from each other). While the first assumption is reasonable, the second one is a simplification, as in practice the dissimilarity between even a fixed pair of haplotypes can vary heavily depending on evolutionary history and chromosomal region. The idea is to estimate the expected Hamming rate between reads from the same haplotype, which we call d_same_, and the expected Hamming rate for reads from different haplotypes, called d_diff_. The former depends only on the sequencing error rate, while the latter additionally includes the differences between the true haplotypes. With d_a_n we further denote the expected Hamming rate over all overlapping read pairs.

For two reads *r* and *s*, the probability of observing the same allele at a shared variant locus equals *d*_same_ if *H*(*r*) = *H*(*s*) or *d*_diff_ if *H*(*r*) ≠ *H*(*s*). Since the variants are independent from each other in our model, dis(r, s) should follow one of the two binomial distributions *B*_olp(*r,s*),*d*_same__ or *B*_olp(*r,*s),*d*_diff__ with olp(*r, s*) being the number of attempts and *d*_same_ or *d*_diff_ being the success probability. For each individual read pair we can then decide which of the two possible distributions is the most likely one.

According to our first assumption, a 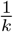-fraction of all possible read pairs include reads from the same haplotype each, as for a read *r* there is a 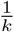 chance that another one is from the same haplotype. In order to estimate *d*_same_ we compute the Hamming rate over all overlapping read pairs and use the average of the lower 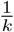-fraction as estimate. As alternative one could also use the sequencing error rate to compute *d*_same_, since the corresponding reads contain only sequencing errors. These error rates are, however, not always available, especially when preprocessing steps like error correction are included. Since *d*_all_ can be simply computed and 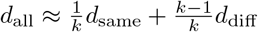, we get an estimate on *d*_diff_ as well. Finally, the similarity score of reads *r* and *s* is defined as

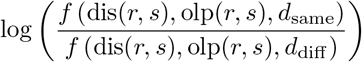

with *f* being the binomial probability density function for dis(*r, s*) successes.

Please note that the score is negative if the read pair appears to be from different haplotypes and positive in the opposite case.

In our studies we noticed that the disagreement rate between haplotypes varied between different regions. In order to increase the accuracy of our model, we partition the variants into windows *w*_1_,…, *w_i_* of average read length and compute *d*_same_ and *d*_diff_ independently for each window. If the overlap region of a read pair spans multiple windows, we use the weighted average of the *d*-values.

As clustering model we chose Cluster Editing [19], which takes a complete graph with real edge weights as input and finds the most cost-efficient way to transform it into a graph only consisting of disjoint cliques. Therefore, positive weighted edges are interpreted as present edges and negative ones as missing edges. The absolute value of a weight is the cost to either insert a missing or delete a present edge. A small example of this model can be found in Figure 5. For our algorithm, we model each read as a node of the input graph and use the similarity score for each read pair to obtain edge weights. Non-overlapping read pairs are defined to have an edge weight of 0, which we call a *zero-edge*. The resulting cliques can be interpreted as clusters of reads with high confidence of originating from the same haplotype.

**Figure 5:**
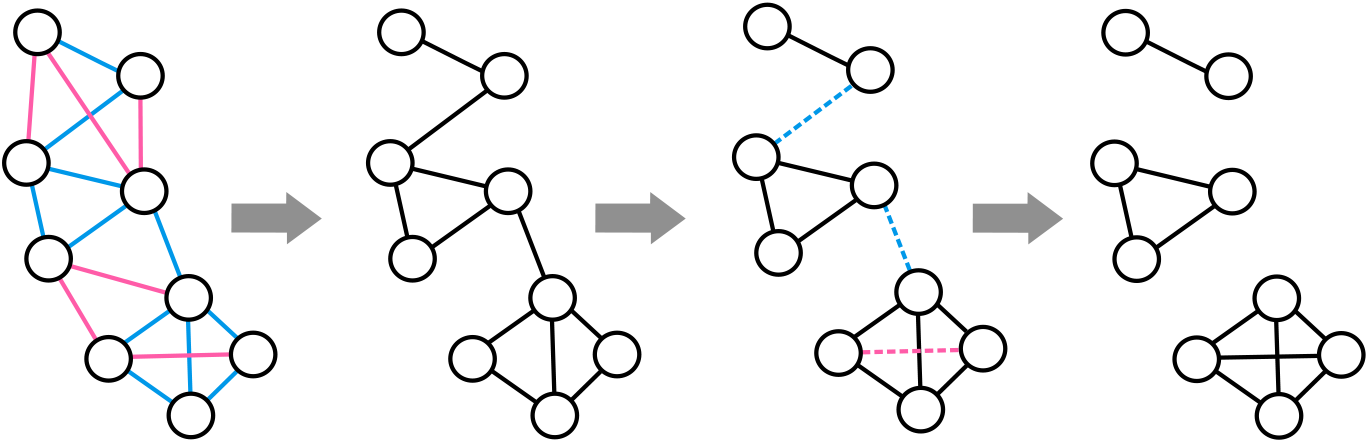
Cluster editing example. The input graph on the left contains one node per read and positive weighted edges (blue) for similar reads and negative weighted edges (pink) for dissimilar reads. All other edges are zero-edges and not drawn for sake of simplicity. The model considers blue edges as present edges and pink edges as missing edges, as shown in the second graph. The information of the pink edges is still used as insertion cost for missing edges. The third graph indicates operations needed to get a clique graph as dashed edges. The blue edges need to be deleted, the pink needs to be inserted. The final clique graph is shown on the right.

The number of clusters depends on the data and is not an input parameter. In practice, we get much more than k clusters for two reasons: First, the distance between variants can vary and can become too large for some variant pairs such that enough reads connect them with sufficient confidence. Second, collapsed regions lead to clusters with reads from multiple haplotypes, forcing single-haplotype clusters to be discontinued. Restricting the cluster editing model to *k* clusters would force clusters to span poorly connected variants and split up reads from locally identical sequences. As this would likely introduce errors, we instead postpone the problem of reducing the clusters to *k* haplotypes to the second part of our algorithm.

Due to the NP-hardness of the Cluster Editing problem, it is infeasible to solve it to optimality on large real-world instances as given by the comparison of all read pairs. Instead, we use a heuristic that greedily picks an edge in each iteration and decides whether it should be present in the resulting clique graph or not, potentially inserting or deleting edges. We denote the first case as making an edge *permanent* and the second one as making an edge *forbidden.* If an edge (*u, v*) is made permanent, for all other nodes *w* it must hold, that either both (*u, w*) and (*v, w*) must be in the final clique graph or none of them. Similarly, if (*u, v*) is made forbidden, there must not be any node *w* such that both (*u, w*) and (*v, w*) are in the final clique graph. Following these conditions, we can compute *induced costs* for each edge (*u, v*), which reflect the costs of obligatory insertion and deletion operations for making (*u, v*) permanent or forbidden. These costs are called icp(*u, v*) and icf(*u, v*) respectively and were originally defined in [31]. Once an edge becomes permanent (forbidden) its weight is set to to (-to) and all induced costs of incident edges are updated accordingly.

To improve the running time, we ignore zero-edges in the heuristic and assume them to not be present in the solution, unless one of them is needed to complete a clique.

### Haplotype Threading

For the second part of the algorithm, we developed a novel approach called *haplotype threading,* which performs the actual phasing to *k* haplotypes. The cluster editing step results in a set *C* of read clusters with two properties: First, the number of clusters at a position *i* ∈ {0,…, *n* – 1} can be larger than *k*, so that some clusters do not contribute to any computed haplotype. Second, the reads in a cluster *c* ∈ *C* usually do not cover the whole chromosome, but only a part of the *n* variants, so in order to obtain whole-chromosome haplotypes, these must be assembled from multiple clusters. This is done by *threading* a haplotype through the clusters, meaning that for every haplotype, a path through *C* is assembled by choosing one cluster *c* ∈ *C* for each haplotype at every variant position *i*.

In a genome of ploidy *k*, we seek for *k* haplotypes and thus assemble all *k* sequences simultaneously by choosing *k*-tuples of clusters at each position. Duplicate clusters within tuples are allowed since reads from one cluster can belong to multiple true haplotypes: For regions with high local similarity between the true haplotypes, the corresponding reads are likely placed into one cluster by the cluster editing step.

In the threading process, we aim at achieving three objectives: (i) genotype concordance, (ii) read coverage and (iii) haplotype contiguity. The first, *genotype concordance*, captures the agreement between the known target genotype and the chosen clusters. For the true haplotypes *H*_0_,…, *H*_*k*–1_ of length *n*, the corresponding genotype can be described as the component-wise sum *G* = *H*_0_ + *H*_1_ + … + *H*_*k*-1_ and is denoted by *G* = *g*_0_, *g*_1_,…, *g*_*n*–1_. Furthermore, for each cluster *c* and each position *i*, we can compute the consensus cons(*c, i*) ∈ {0,1, −} as the most frequent allele among all reads in *c* at position *i*. Using this definition, we can compute a consensus genotype of a *k*-tuple (*c*_0_,…, *c*_*k*–1_) at position *i* as 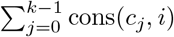. For each position *i*, we then only take those cluster tuples into account whose consensus genotype at *i* is *concordant* with the target genotype, i.e., 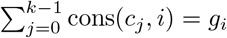. Reducing the search space of possible tuples this way increases efficiency as well as accuracy by filtering out non-promising combinations beforehand. In case there is no tuple with a concordant genotype at position *i*, we allow genotype deviations of 1; if this also fails, all possible tuples are considered.

To determine the best fit among the possible cluster tuples, we designed an objective function that takes the remaining two criteria into account as follows. The second criterion is *read coverage.* Since in locally identical regions multiple haplotypes can be threaded through the same cluster—which leads to multiple appearances of this cluster in the *k*-tuple—this number of haplotypes has to correspond to the coverage of the chosen cluster. The relative coverage of a cluster *c* at position *i* describes the number of reads in *c* covering *i* divided by the total number of reads in all clusters that cover *i*. We denote this value by cov(*c, i*). Then, we can compute the expected copy number of a cluster *c* at *i*, i.e. the expected number of haplotypes that are threaded through *c*, by 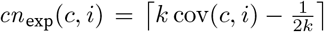. The true copy number of *c* in a chosen cluster tuple (*c*_0_,…, *c*_*k*–1_) is given by *cn*_true_((*c*_0_,…, *c*_*k*–1_),*c, i*) = |{*i*| *i* ∈ {0, …, *k* – 1}, *c* = *c_i_*}|. Deviations of the true number of occurrences from the expected ones are penalized by a constant factor *p*_cov_ per cluster, so that a cluster tuple (*c*_0_,…, *c*_*k*–1_) is evaluated by the cost function

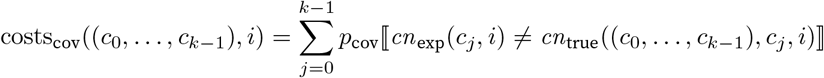

where [*x* ≠ *y*] returns 1 if *x* ≠ *y* and 0 otherwise.

The third and last criterion, *haplotype contiguity,* encourages haplotypes to stay in the same cluster as long as possible, so that switching of haplotypes between clusters is penalized. For two consecutive cluster tuples (*c*_0_,…, *c*_*k*–1_) and 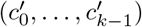 at positions *i* and *i* + 1, we denote the cost factor by *p*_switch_, which results in the cost function

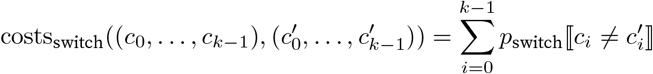

We developed a dynamic programming approach to rapidly find the optimal sequence of tuples that minimizes all costs. We compute a two-dimensional matrix *S* with a column for every variant *j* from 0 to *n* – 1 and a row for every possible genotype-conform tuple of clusters. Since the number of eligible cluster tuples can differ between variant positions, the columns of S do not necessarily have the same lengths. We denote the length of a column *j* with *l_j_*. Using the cost functions defined above, *S*[*i, j*] is then computed as

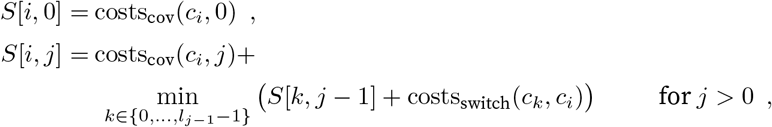

where *c_i_* denotes the cluster tuple in row *i*. The optimal threading score is then given by the minimum value in the last column. Starting at this position, we assemble the sequence of clusters with minimum costs via backtracing.

The threading process is illustrated in Figure 6a for *k* = 4. The clusters from the first step are drawn as grey shapes in a two-dimensional space, where the horizontal position refers to the variants covered by the reads inside a cluster and the height represents the relative coverage of a cluster at every position. The position on the *y*-axis has no numerical meaning and is just used for illustration purpose. Starting from the left, a 4-tuple of the five present clusters needs to be chosen. According to the coverage, the best choice is to thread one haplotype through each of the four clusters with highest coverage and ignoring the smallest one, as this is likely to contain noisy reads only. From thereon, the threads change clusters whenever a cluster ends or undergoes a drastic change in relative coverage.

**Figure 6:**
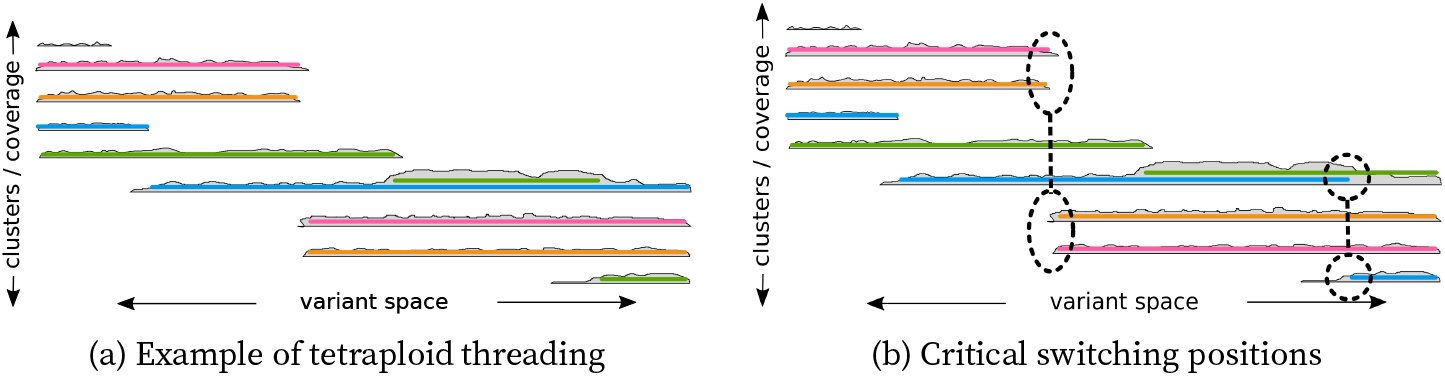
Visualization of the threading. (a) Clusters of reads are represented as grey shapes with their horizontal span indicating the covered variants and the height being the respective coverage. The *k* = 4 threads are shown as colored lines passing through the clusters. Multiple threads can co-enter the same cluster if the coverage is suited. (b) Alternative threading with the same score in our model. Two positions cause ambiguity and allow switches in the threading compared to (a). These are candidate cut positions to prevent switch errors in the final phasing.

### Block cuts

Phasing tools are able to divide phased haplotypes into blocks if there is not enough evidence in the data to connect these blocks. This is usually done when there are two variants with no connecting read in between. For polyploid organisms, however, even a single connecting read is not sufficient, as reads from *k* – 1 different haplotypes are needed to resolve the connection of *k* haplotypes on both sides. In general, block cuts are a trade-off between block length and accuracy, as one of these metrics can easily be optimized by giving up the other one. To offer more flexibility to the user, WhatsHap polyphase provides different modi of applying block cuts to either get short and accurate or long but less accurate blocks.

Since it is uncertain whether different read clusters represent the same or different haplotypes, the most conservative method is to cut the phasing whenever one thread switches to another cluster, which we call *single-switch-cuts.* While this yields the lowest block-wise error rate, many of these cuts can be avoided. If only one thread switches the cluster, while the other *k* – 1 threads stay, one could conclude that the old cluster is linked to the new one by process of elimination. If two or more threads switch, the continuation is ambiguous and a cut can be placed here to prevent switch errors, which we call a *multi-switch-cut*. In principle, only the switching threads need to be cut, while the rest can stay connected. To the best of our knowledge, however, there is no established method to express such selective block cuts in a VCF file. Therefore, all haplotypes are interrupted in case of a multi-switch-cut.

The last type of cuts is the *separation-cut*, which is necessary to handle collapsed regions. Assume a cluster contains multiple threads at some position and the number of threads has to be decreased by 1 for the next position due to a decrease in coverage. Even though this is not covered by the multi-switch-cuts, there is still a choice which of the contained threads should leave the cluster. If all threads have been in the cluster since the start of the current block, the leaving thread can be chosen arbitrarily. However, if they entered the current cluster on different positions or from different predecessor clusters, the choice affects the resulting haplotype sequences and we need to insert a separation-cut here to avoid potential switch errors. Figure 6b shows an example, where two threads (green and blue) share the same cluster before one of them has to leave. Either of them switching would lead to a different result, for which we do not know the correct one.

### Preprocessing and phasing the potato genome

We ran a recently developed error correction pipeline [25] to reduce the typically high number of sequencing errors in the Oxford Nanopore MinION reads, in order to use them for variant calling and phasing. Illumina reads were first self-corrected using Lighter [32], the corrected reads were used to build a de Bruijn graph with bcalm2 [33] and the MinION reads were aligned to the graph with GraphAligner [25]. We used the default parameters for the error correction pipeline (Lighter *k* = 21, bcalm2 *k* = 61 and abundance = 3, GraphAligner default alignment parameters). The corrected read sequence of each mapped MinION read was obtained from the path of its respective alignment in the graph. The corrected reads where aligned to the reference genome using minimap2 [26] and converted to BAM-format using samtools [34]. In the next step, we ran FreeBayes [27] (with additional parameters: `-p 4 -no-indels -no-mnsp -no-complex`) inside of all gene regions to call SNPs from the corrected Nanopore alignments. As base qualities are not produced during error correction and FreeBayes seems to need them in order to compute genotypes, we added a constant quality of 40 for all bases to the BAM file before calling SNPs. Finally, we ran WhatsHap polyphase in order to phase the variants with option `-verify-genotypes`. This option invokes an additional step prior to phasing, which re-genotypes all variants and only keeps those positions for which the computed genotype matches the input genotype. For determining the genotype of a position, we implemented a simple algorithm that calculates the fraction of reference and alternative alleles that cover a variant and compare it to the fractions that we would expect for all possible genotypes. We then assign the genotype whose expected fractions of reference and alternative alleles best match the ones observed in the data.

We focused on the FRIGIDA-like protein 5 isoform X2 gene to demonstrate a use case of polyploid phasing. We first extracted all phased variants that are part of the longest phasing block reported by WhatsHap polyphase for this gene. In order to assign reads to the haplotypes computed by WhatsHap polyphase, we extended the command `whatshap haplotag`, which was previously implemented for the diploid version of `whatshap`, to the polyploid case. Given a phased VCF with predicted haplotypes and BAM-file with sequencing reads, we assign each read to the haplotype it is most similar to in terms of the alleles observed at variant positions in the read. This assignment is stored by tagging the respective sequences in the BAM file, which enables visualizing the haplotype clusters by programs like IGV [23] (see Figure 4c). Furthermore, we extended the subcommand whatshap split to higher ploidies, which can be used to split tagged reads by haplotype and store them in separate files. For each haplotype, we produced a BAM-file with reads in this way.

In the next step, we ran wtdbg2 [28] (with options `-x ccs -g 1m`) separately for reads corresponding to each haplotype to generate haplotype-resolved assemblies for the Frigida gene. Those were further analyzed with NCBI’s ORFfinder and COBALT algorithms [29, 30] using their web interfaces (https://www.ncbi.nlm.nih.gov/orffinder/, https://www.ncbi.nlm.nih.gov/tools/cobalt/re_cobalt.cgi).

## Availability of data and materials

WhatsHap polyphase is available at https://bitbucket.org/whatshap/whatshap. All scripts needed to generate data and reproduce the analysis can be found at https://github.com/eblerjana/whatshap-polyphase-experiments. Sequencing datahas been submitted to the NCBI with accession number PRJNA587397.

## Acknowledgements

Thanks to Sebastian Böcker for the idea to use the icp and icf values in a greedy cluster editing heuristic.

## Supplement

### Supplement 1 — Hamming rates for artificial tetraploid human

**Figure 7:**
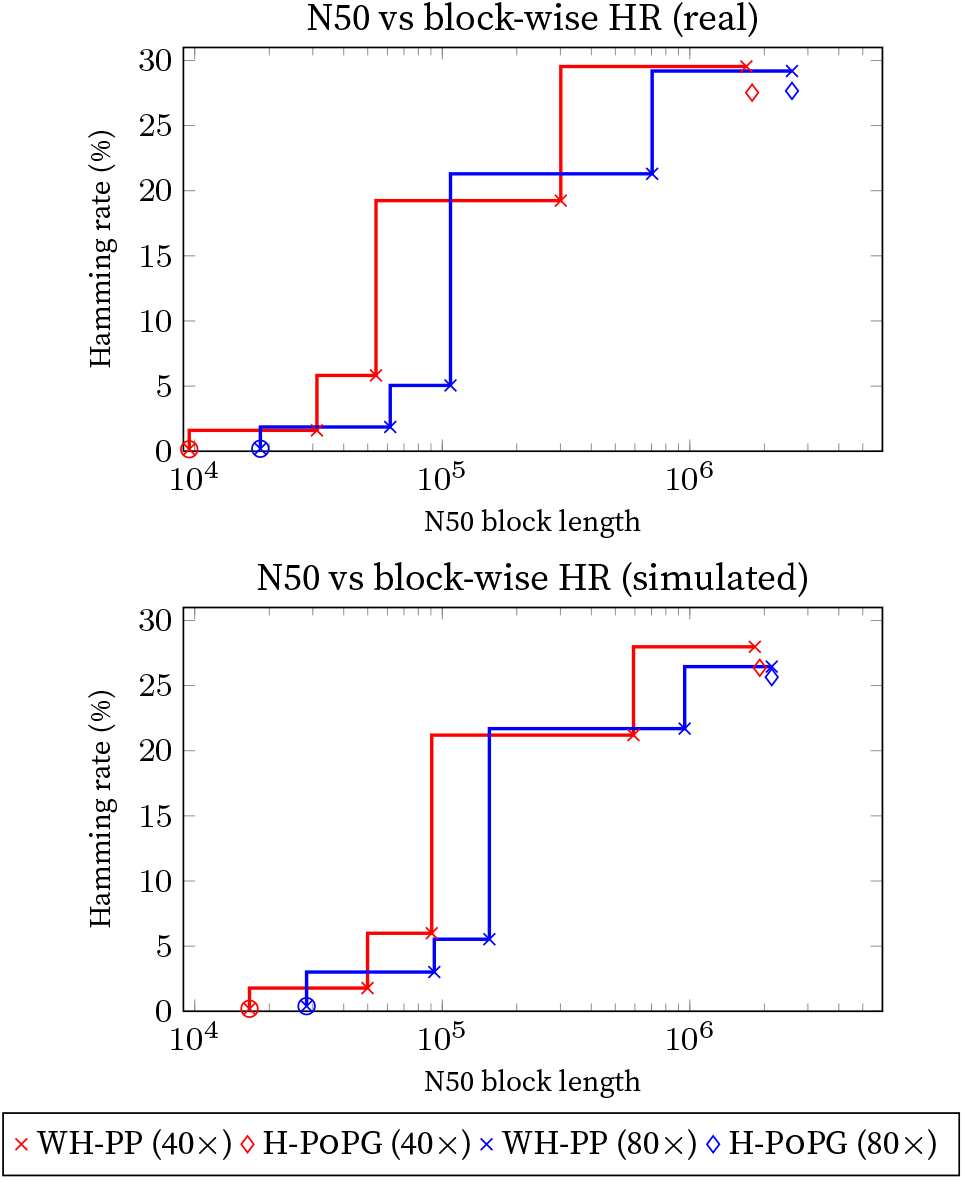
N50 block lengths and the respective block-wise Hamming rates for different block cut strategies of WhatsHap polyphase (default strategy marked by a circle) on the real read dataset (top) and the simulated dataset (bottom) with 40 × and 80 × coverage. Note that Hamming rates above 10 or 20 percent do not seem to be useful in practice. If one used such a phasing to query whether two alleles lie on the same haplotype, the chance of error would be as high. The results illustrate that both methods are unable to produce good phasings over longs blocks. WhatsHap polyphase achieves reasonably low Hamming error rates the default block cut strategy.

### Supplement 2 — Comparison between H-PoPG and WhatsHap polyphase in collapsing regions, using long blocks similar to H-PoPG

**Table 3:** Comparison between the resulting switch error rates of H-PoPG and WhatsHap polyphase using block lengths that are comparable to H-PoPG (WH-PP-H) on collapsing regions over at least 50 variants as compared to non-collapsing regions and the average throughout the genome. Results (switch error rates in %) are presented for Chr01 of the real (a) and the simulated (b) dataset, testing 40 × and 80× coverage. The third row marks the quotient between the switch error rate of H-PoPG and that of WhatsHap polyphase to highlight by which magnitude the results differ.

| coverage | method | collapsing regions | non-collapsing regions | total |
| --- | --- | --- | --- | --- |
| 40× | WH-PP-H | 0.66 | 1.81 | 1.65 |
|  | H-POPG | 2.02 | 2.16 | 2.02 |
| | SER($\frac{H-POPG}{WH-PP}$) | 3.06 | 1.19 | 1.22 |
| 80× | WH-PP-H | 0.38 | 1.16 | 0.99 |
|  | H-POPG | 1.05 | 1.30 | 1.24 |
| | SER($\frac{H-POPG}{WH-PP}$) | 2.76 | 1.12 | 1.25 |

(a) real read data
| coverage | method | collapsing regions | non-collapsing regions | total |
| --- | --- | --- | --- | --- |
| 40× | WH-PP-H | 0.45 | 1.29 | 1.19 |
|  | H-POPG | 2.01 | 1.63 | 1.68 |
| | SER($\frac{H-POPG}{WH-PP}$) | 4.47 | 1.62 | 1.41 |
| 80× | WH-PP-H | 0.25 | 0.88 | 0.82 |
|  | H-POPG | 0.94 | 0.98 | 0.99 |
| | SER($\frac{H-POPG}{WH-PP}$) | 3.76 | 1.11 | 1.21 |

(b) simulated read data

### Supplement 3 - Alignment of the haplotype sequences of the FRIGIDA gene and the reference genome

**Figure 8:**
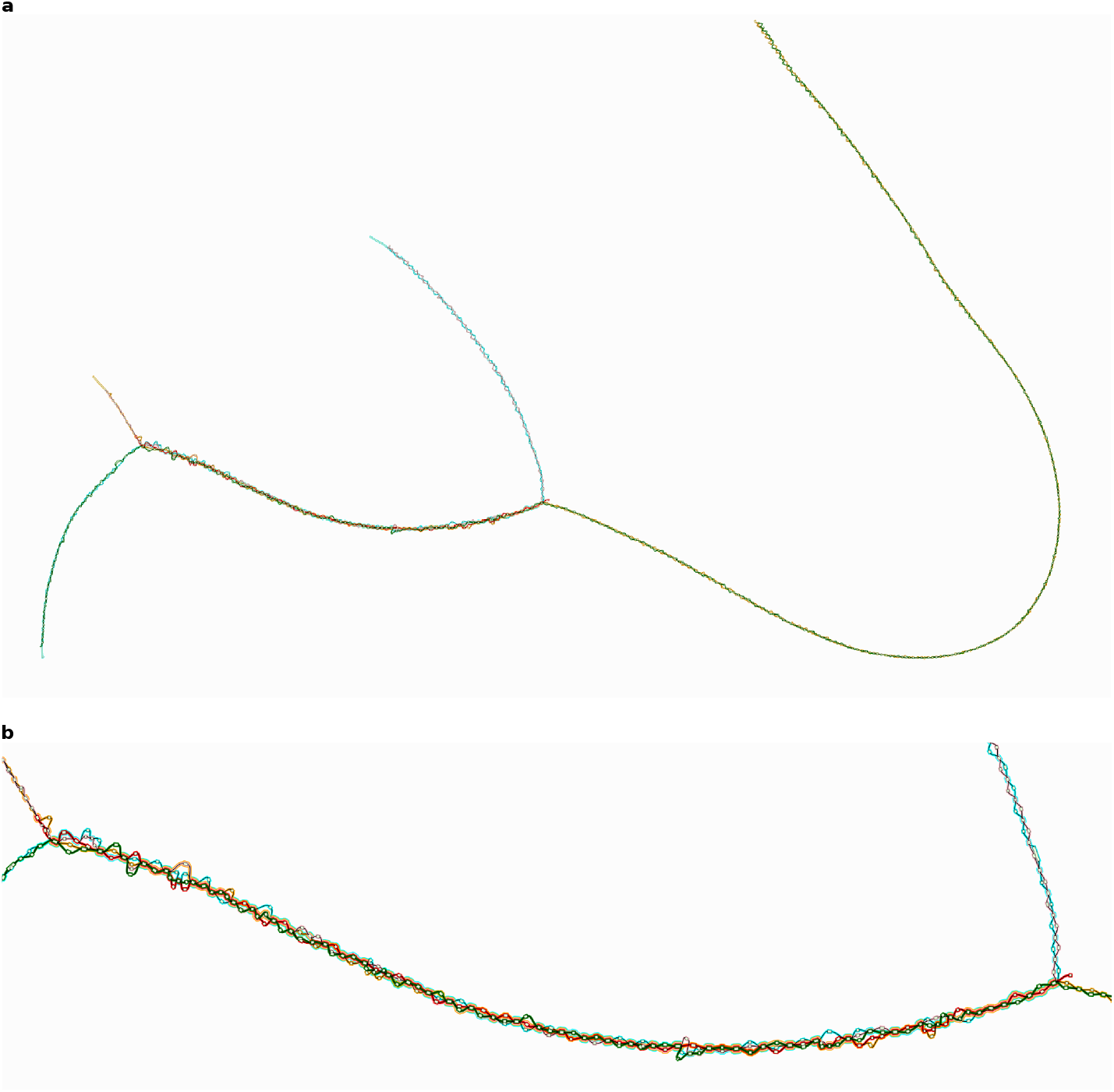
Haplotype assemblies for the FRIGIDA gene. We ran Reveal https://github.com/jasperlinthorst/reveal to produce a graph that represents an alignment of the local haplotype assemblies for the FRIGIDA gene and the corresponding reference sequence. We visualized this graph using GFAviz [35]. The red sequence corresponds to the reference genome. **a)** shows the whole graph, **b)** shows the part of the graph that corresponds to the FRIGIDA-like protein 5 isoform X2 gene.

